# Design and Validation of a Frugal, Automated, Solid-Phase Peptide Synthesizer

**DOI:** 10.1101/2020.05.21.109215

**Authors:** Nathaniel E. Kallmyer, Nathan E. Rider, Nigel F. Reuel

**Author notes:** Corresponding Author –.

## Abstract

Solid phase peptide synthesis (SPPS) has enabled widespread use of synthetic peptides in applications ranging from pharmaceuticals to materials science. The demand for synthetic peptides has driven recent efforts to produce automated SPPS synthesizers which utilize fluid-handling components common to chemistry laboratories to drive costs down to several thousand dollars. Herein, we describe the design and validation of a more ‘frugal’ SPPS synthesizer that uses inexpensive, consumer-grade fluid-handling components to achieve a prototype price point between US$300 and $600. We demonstrated functionality by preparing and characterizing peptides with a variety of distinct properties including binding functionality, nanoscale self-assembly, and oxidation-induced fluorescence. This system yielded micromoles of peptide at a cost of approximately $1/residue, a cost which may be further reduced by optimization and bulk purchasing.

## Introduction

Synthetic peptides are a topic of increasing study for many applications that range from drug delivery to novel materials and sensing components[1,2]. These peptides are commonly produced by solid-phase peptide synthesis (SPPS), a foundational approach for which Bruce Merrifield won the Nobel Prize in Chemistry, where amino acids are consecutively added to a solid resin by amidation to effectively grow peptides one residue at a time[3]. SPPS involves a repetitive strategy in which protected amino acids are added, and then a protection group is removed to allow addition of the next residue. The ability to synthesize peptides by SPPS has greatly improved as new linking reagents and protection groups have allowed use of increasingly modular and decreasingly hazardous reagents[4,5]. Fragment condensation and post-translational modification techniques have enabled synthesis of molecules resembling natural proteins[6–9]. Combinatorial prototyping strategies have also allowed preparation of large libraries in single batch reactions[10]. Improvements to synthesizers, including continuous-flow and microwave-assisted techniques, have allowed ultra-rapid fabrication for high-throughput prototyping synthesis[11–16]. While these new tools have proven effective, they require expensive equipment or specialized fabrication tools. The simplicity and repetitiveness of SPPS presents an opportunity to develop a minimalist tool which can bring synthetic peptides to any research group at an accessible price.

The development of “frugal-science” low-cost tools to address large-volume problems or challenges in third-world countries has been an increasing area of study[17]. To address financial limitations, there has been a push to develop tools that use simple designs and inexpensive components, such as the paper centrifuge or origami microscope[18,19]. Similarly motivated, open-source designs have been applied to SPPS; however, these designs still rely on expensive fluid-handling components adapted from chromatography apparatuses, raising prices to the range of several thousand USD[13,20]. By replacing fluid-handling equipment with consumer-grade components, such as aquarium pumps, it is possible to substantially reduce synthesizer build costs. Herein, we demonstrate a low-cost solid phase peptide synthesizer that can be manufactured for under US$300 using inexpensive peristaltic pumps, coupled with electrical relays and a microcontroller.

## Methods

### Synthesizer Construction

The peptide synthesizer reactor comprised of two 10-mL columns connected serially. The first column was used to premix and activate fmoc-amino acids. These activated amino acids were then pumped into the second column for amidation of the resin. Fluids were delivered by peristaltic pumps (Yosoo) through silicone tubing. The peptide synthesizer was controlled by a Raspberry Pi 3B which operated two 8-channel relays (Sunfounder) that triggered peristaltic feed pumps powered by a 12 V DC power supply. Wires from the power supply were soldered to the relay and peristaltic pump inputs and shrink wrapped. A cylindrical neodymium magnet was fixed to the shaft of a small DC motor (Pimoroni COM0805), powered by a 7.5 V DC power supply, and used to drive a magnetic stirrer. A wiring diagram of the pump control components is provided in Supplement 1.

### Peptide Synthesis

DMF and piperidine were obtained from Millipore Sigma. Protected amino acids, resins, and amidation reagents were obtained from Chem-Impex. A comprehensive list of reagents is provided in Supplement 2. Reagent preparation and peptide synthesis was performed inside a fume hood. Peristaltic pumps were used to perform six different functions. One of the pumps fed pure dimethylformamide (DMF) to rinse resin between steps. A second pump fed 20% v/v piperidine in DMF for deprotection of amino acids. A third pump transferred activated reagents from an upper premixing vessel to the bottom, resin-containing reaction vessel. A fourth pump transferred liquid from the bottom reaction vessel to a waste container. The fifth pump fed an amino acid activation solution (75 mg/mL 1-[Bis(dimethylamino)methylene]-1H-1,2,3-triazolo[4,5-b]pyridinium 3-oxide hexafluorophosphate, Hexafluorophosphate Azabenzotriazole Tetramethyl Uronium (HATU) in 4.3% v/v N-methylmorpholine (NMM)/DMF) into the upper pre-mixing vessel. The sixth and remaining pumps fed different fmoc-protected amino acids (0.1 M fmoc-amino acid in DMF) into the premixing vessel for activation.

The SPPS process is outlined in Fig. 1. Prior to SPPS, all tubing was flushed with DMF and primed. To begin, 50 mg of resin (aminomethyl polystyrene for bead assays, chlorotrityl resin for free peptide tests) were loaded into the reaction vessel (bottom vessel) with approximately 2.5 mL of DMF. Next, the resin was swelled for approximately 15 min. A python script, provided in Supplement 3, was then initiated to perform the remaining steps (Fig 1). First, the bottom (resin-containing) vessel was drained by peristaltic pump. Then, two 0.75- mL rinses of 20% v/v piperidine were added to this bottom vessel (5 min for each). Next, the reaction vessel was rinsed 5 times each with 2 mL DMF. After rinsing, 1.25 mL of the fmoc-amino acid solution and 0.625 mL of the activation solution were added to the premixing vessel. After 10 s, the contents of this vessel were then drained into the reaction vessel. 15 min were given for the amidation reaction to occur. Following amidation, the premixing vessel was rinsed three times and drained. The resin in the reaction vessel was then rinsed with 20% piperidine, and this procedure was repeated for each amino acid. Throughout this procedure, the reaction vessel was agitated by a micro-stir bar at 400 RPM. Following peptide growth, 2 mL 90% trifluoroacetic acetic acid in DMF were added to the reaction vessel to remove protection groups from peptide side-chains and, in the case of chlorotrityl resin, cleave the peptide from the resin. Following completion of SPPS, silicone tubing was rinsed with deionized water to minimize degradation of tubing by DMF between synthesis runs. Sequences of synthesized peptides are provided in Table 1. An additional glycine was included on the fluorescent peptide to allow synthesis on a glycine-functionalized chlorotrityl resin.

**Figure 1.**
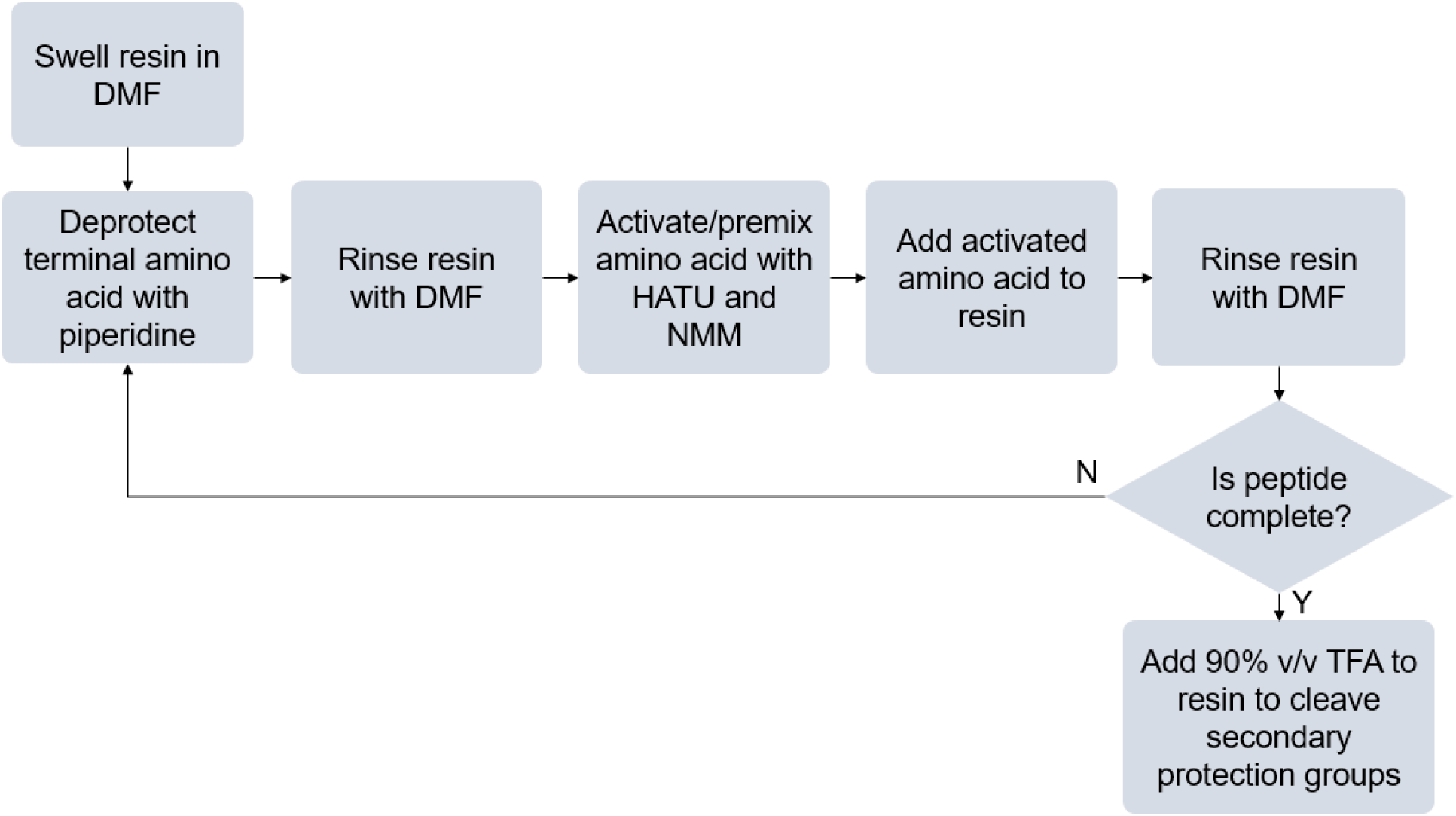
Process flow diagram of SPPS process, beginning with swelling of resin and ending with removal of side-chain protection groups.

**Table 1.**
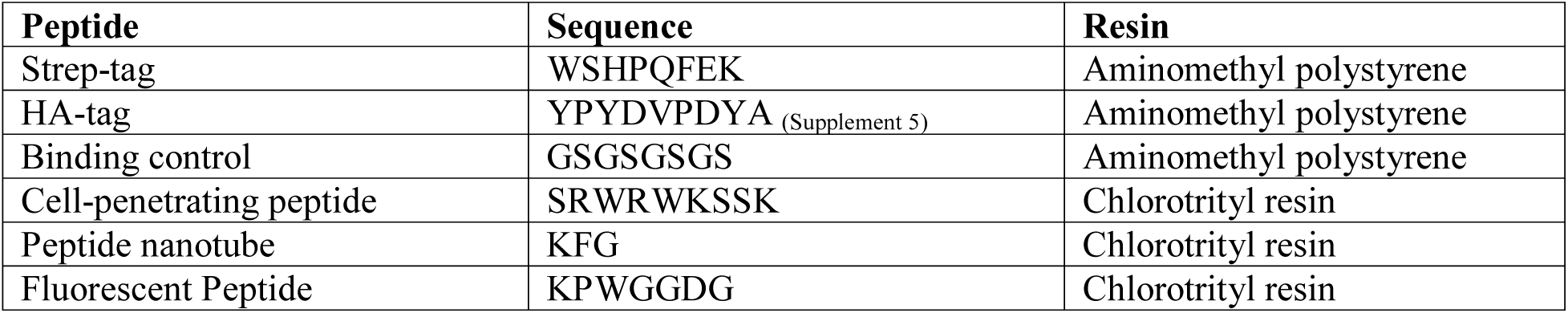
Sequences of Peptides Synthesized with Frugal System.

| Peptide | Sequence | Resin |
| --- | --- | --- |
| Strep-tag | WSHPQFEK | Aminomethyl polystyrene |
| HA-tag | YPYDVPDYA (Supplement 5) | Aminomethyl polystyrene |
| Binding control | GSGSGSGS | Aminomethyl polystyrene |
| Cell-penetrating peptide | SRWRWKSSK | Chlorotrityl resin |
| Peptide nanotube | KFG | Chlorotrityl resin |
| Fluorescent Peptide | KPWGGDG | Chlorotrityl resin |

### Binding Affinity Tests

Aminomethyl polystyrene resins were then rinsed repeatedly with 2.5-mL additions of deionized water (appr. 20 rinses) until the pH rose above 6. For binding assays, resins were first incubated in fluorophore-tagged antibodies (FITC-streptavidin, Vector Labs SA-1200) for 20 minutes. Following incubation, the beads were rinsed with 10 additions of 2.5 mL deionized water. These beads were then compared with a fluorescent microscope at identical exposure, gain, and contrast settings. Beads were selected by bright-field spectroscopy prior to measurement of fluorescence to avoid any bias when measuring bead emission.

### Free Peptide Characterization

Peptides produced on chlorotrityl resin were cleaved by trifluoroacetic acid and collected as effluent. The resin was then rinsed with 1.5 mL isosopropyl alcohol to collect any residual peptide. The trifluoroacetic acid and isopropyl alcohol were then evaporated with an in-house dry air line. Next, 2 mL supercooled (−80 ºC) diethyl ether was added to the peptide-DMF solution to precipitate peptide product. The DMF phase was then extracted, and the diethyl ether was evaporated to obtain solid peptide product.

Peptide yields were predicted by absorbance spectroscopy. Yields of the cell-penetrating peptide were determined by comparing tryptophan absorbance at 280 nm to that of a peptide standard (Genscript). This peptide product was verified by time-of-flight electrospray ionization cationic mass spectroscopy Agilent QTOF 6540). Concentrations of nanotube-forming peptide (KFG) were determined by comparing phenylalanine absorbance at 257 nm to a phenylalanine standard. After neutralizing residual trifluoroacetic acid with 0.5 M NaOH, 0.5 mg/mL and 2.5 mg/mL KFG peptide solutions were spotted onto cleaved mica slides. After the solutions dried, topography was scanned by non-contact mode atomic force microscopy (Agilent 5400).

### Fluorogenic Peptides

Following synthesis, cleavage from the resin, and purification, the fluorogenic peptide was stored in a 5-mL centrifuge tube, and the tube was wrapped with aluminum foil to minimize further exposure to light. This peptide was then dissolved in 1 mL deionized water and separated into 2 500-µL aliquots, one control and one sample to be photooxidized. The latter was affixed to a halogen lamp and illuminated at 240 W at 5 min intervals. Following each interval, an 80 µL sample was collected and saved for fluorescent scanning. Samples were loaded onto a 96-well plate, and fluorescence was scanned (ex = 340 nm, em = 393 nm) by a microplate reader (Biotek Synergy Neo2). Four analytical replicates were collected.

### Results and Discussion

The repetitive steps of SPPS allows for construction of a minimalist reactor system consisting of only two reaction vessels: a pre-mixing vessel and amidation vessel (Fig 2). By substituting relatively expensive piston pumps with peristaltic pumps, typically used in aquariums, a larger number of inexpensive pumps may be used as a substitute for valves and valve actuators. Removal of shared fluid lines also minimizes potential sources of contamination. If finer fluid control is required, peristaltic pump motors can be substituted with higher torque, more easily controlled motors. To reduce synthesis time, reagents can be preheated in an incubator or in the feed line by Joule heating. More feed lines can be added easily by attaching additional peristaltic pumps. The number of Raspberry Pi outputs can be substantially increased to accommodate these additional pumps by inclusion of a binary encoder.

**Figure 2.**
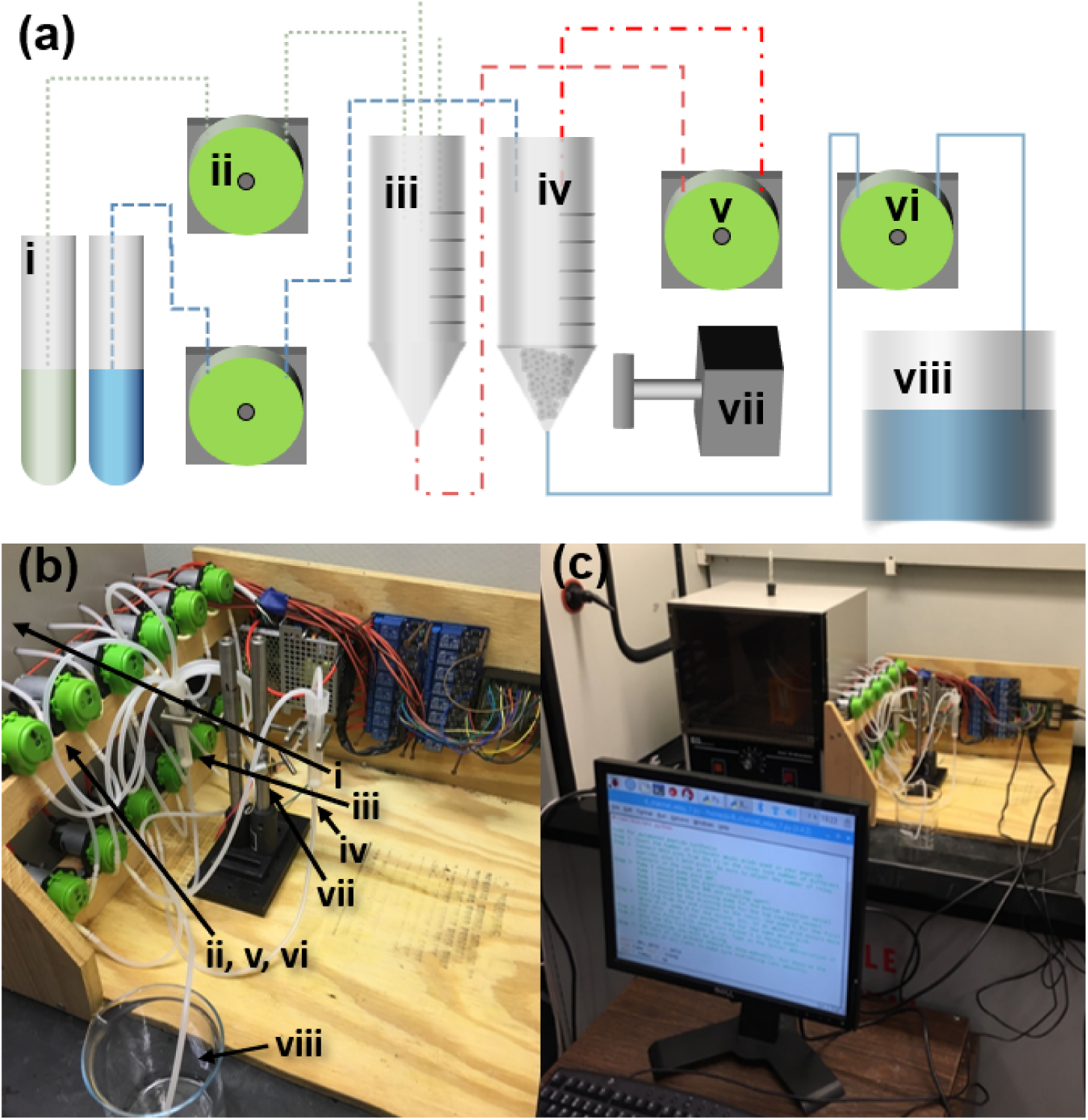
Design of solid phase peptide synthesizer fluid-handling components. **(a)** Schematic of fluid-handling components. (**i**) Reagents for SPPS, (**ii**) reagent feed pumps, (**iii**) reagent mixing and activation vessel, (**iv**) resin and peptide synthesis vessel, (**v**) transfer pump from activation to peptide synthesis vessel, (**vi**) drainage pump from, (**vii**) custom, inexpensive magnetic stirrer, and (**viii**) waste container for reactor effluents. **(b)** Photograph of fluid handling components. **(c)** Photograph of complete setup and physical interface.

The low cost of the peptide synthesizer is enabled by the simple design and inexpensive fluid handling components. This prototype, capable of handling 6 amino acids, cost approximately US$300 (Table 2). With additional pumps and control elements, this design could be used to simultaneously handle all 26 natural amino acids while keeping the total price below US$600.

**Table 2.** Parts List and Itemized Cost.

| Part | Quantity/Length | Supplier | Price (US\$) |
| --- | --- | --- | --- |
| Raspberry Pi 3b+ | 1 | Digikey | 40 |
| Silicone tubing 6mm x 8 mm | 2m | uxcell | 12 |
| 8-channel relay module | 2 | Sunfounder | 24 |
| Peristaltic Aquarium Pump | 12+ | Yosoo | 144 |
| Pierce Disposable 10 mL column | 2 | Thermo Fisher | 5* |
| 12 V 25 W Converter | 1 | Digikey | 23 |
| 22-gauge electrical wire | 2m | BNTECHGO | 3* |
| 3-connector electrical cable | 2 | Digikey | 8 |
| Neodymium cylindrical magnet | 1 | Digikey | 2 |
| 7.5 V 41 W Converter | 1 | Digikey | 18 |
| Micro metal gearmotor | 1 | Digikey | 8 |
| **Total** |  |  | 287 |

Yields of the cell penetrating and nanostructured peptides, determined based on absorbance at 280 nm, were 2.47 mg (1.84 μmol) and and 2.17 mg (6.18 μmol) respectively. Based on these yields, this process utilized reagents (amino acids and HATU) at a 1-5% efficiency. This low efficiency was by design, as an excess of activated amino acids was used to maximize rate of reaction. While this efficiency may be improved, it would require optimization of reaction times and reagent concentrations with consideration of ambient temperature. Even at a single percentage efficiency, this synthesizer produced micromoles of peptide at a cost of approximately US$1/residue. This price was calculated based on reagents purchased for this study. Calculations are provided in Supplement 5. With further price comparison and bulk purchasing, it would be possible to reduce this reagent price.

### Safety Considerations

In the development of new, low-cost tools, it is necessary to address the safety constraints of this process. Although the low-cost of this prototype makes SPPS accessible to a wide user base, the chemicals employed in this process require precautions that should restrict use. Due to hazardous and volatile reagents, this system was run in a fume hood and was only operated by individuals trained in laboratory safety procedures. Personal protective equipment included nitrile gloves, polycarbonate safety glasses, lab coats, pants, and closed toed shoes. Solvents and reagents were stored in a flammables cabinet when not in use.

Because this system handled flammable, organic solvents, prevention of contact between flammable solvents and electrical components was critical. Pumps, relays, and the Raspberry Pi computer were elevated to prevent accumulation of fluid on electrical components in the case of a leak. All connection and exposed wires were insulated by shrink wrap. The prototype employed in this work was monitored (not left unattended), and an ABC fire extinguisher was kept in the laboratory in case of any equipment malfunction and fire. DMF, the solvent used in this process, in addition to being flammable, is toxic, carcinogenic, and easily absorbed through skin. Piperidine, TFA, and NMM are also toxic and corrosive. Waste should be stored in a closed container and sent to a waste management facility.

### Synthesizer Performance

Due to the automation of SPPS, most operator hours were spent preparing stock solutions and flushing silicone tubing. One distinguishing feature of this frugal synthesizer was the use of a magnetic stir bar for agitation rather than compressed gas. While this introduced an issue related to resin damage and nonspecific binding to the fractured, newly exposed resin, these damaging effects are mitigated by reducing size of the stirrer and rate of rotation. Use of a stirrer allowed a simpler reactor design, eliminating the need for a shared drainage and aeration line. Removal of this shared line also reduced risk of reactor contamination by backflow of fluid in the line. One concern about this design was the lifetime of the silicone tubing with the DMF solvent. When flushed with water after use, silicone tubing showed no noticeable damage after over 80 hours of synthesis used in this project.

### Characterization of Affinity Tag Peptides

The resulting resin with synthesized strep-tag exhibited a clear binding preference to fluorescently labeled streptavidin relative to (GS)_4_ controls (Fig 3). Fluorescence observed on the controls was associated with non-specific binding of streptadivin to unprotected (fractured) polystyrene. Non-specific binding was most pronounced on the damaged or fragment resin beads. This resin damage was associated with use of a stir bar at a non-optimal rate, rather than bubbling nitrogen through the solution. Pixel intensities were quantitively compared by selecting regions of interest between Strep-tag beads and control beads. A similar test was performed on beads with HA-tags made by the frugal synthesizer (Supplement 5); however, intensity trends were not as clear as the stronger Strep-tag.

**Figure 3.**
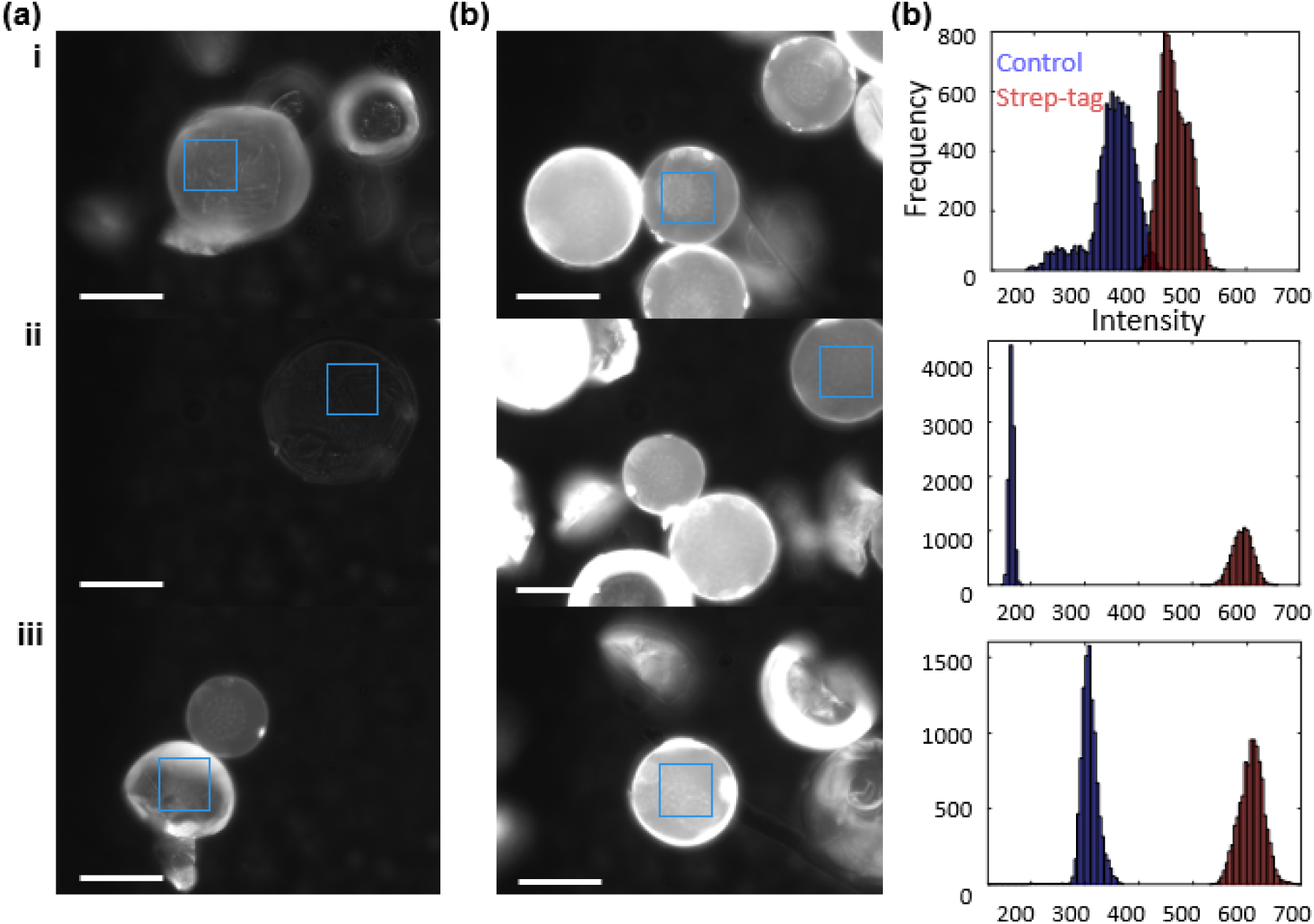
Fluorescent microscope image of aminomethyl styrene resin functionalized with **(a)** control (GS)_4_ sequence and **(b)** strep-tag after incubated with FITC-streptavidin. **(c)** Histogram comparison of pixel intensities collected from regions of interest (solid blue boxes in a-b). **(i-iii)** Replicates. Scale bars are 50 µm.

### Characterization of Cell-penetrating Peptide

A common use of solid-phase peptide synthesis is prototyping therapeutic agents, often small amphiphilic molecules that may function as tags to deliver biopharmaceutical cargo to the cell cytoplasm. Thus, a cell-penetrating peptide based on Cylop-1 was selected for testing on this frugal system[21,22]. Following synthesis, this peptide was characterized by electrospray ionization mass spectrometry (Fig 4). A peak at m/z = 1348 (z = 1) was expected when testing this peptide, but no single-charge peak was visible in either commercially synthesized standard or frugally synthesized sample. Peaks indicated multiply charged cations (+3, +4, +5, +6) were abundant (m/z = 450, 337, 271, 192). The prevalence of multi-charged cations may be explained by the large quantity of basic residues (lysine, arginine, tryptophan) in the sequence. Additionally, multicharged peaks were more prominent in the frugally synthesized product. It is possible that this disparity results from a lower pH of the synthetic product due to residual trifluoracetic acid.

**Figure 4.**
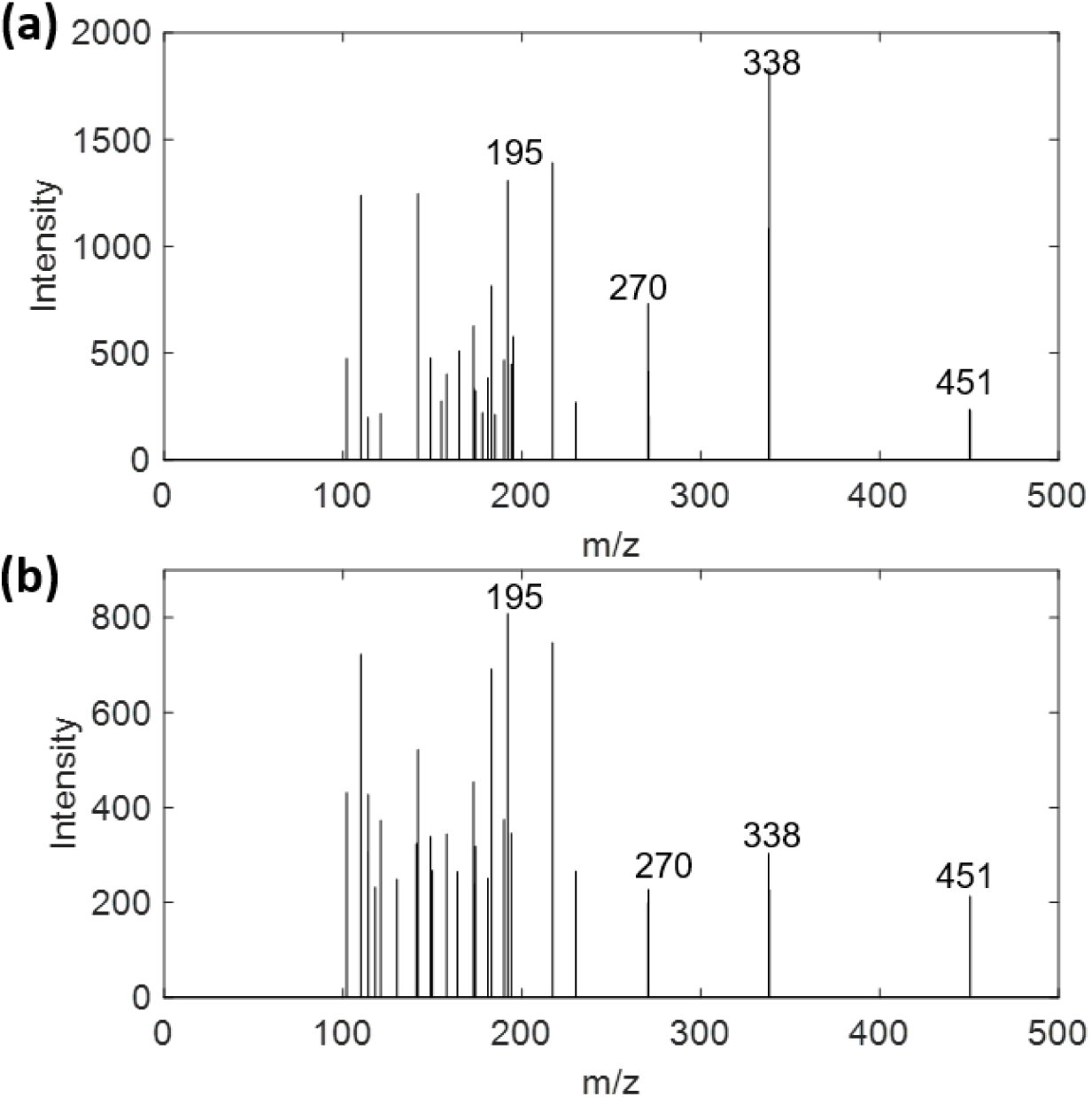
**(a)** Electrospray ionization mass spectrometry cation spectrum of cell-penetrating peptide standard (m = 1347). **(b)** Mass spectrum of frugally synthesized cell-penetrating peptide. Multiply charged peaks associated with peptide are labeled with m/z value.

### Characterization of Nanostructured Peptide

Peptides can also be synthesized to produce self-assembling nanostructures with potential utility in novel materials and pharmaceuticals. For this study, a nanotube-forming tripeptide, KFG, demonstrated as a self-assembling monomer by Moitra et. al., was synthesized and scanned by AFM[23]. At the low peptide concentration of 0.5 mg/mL, small agglomerate structures were produced (Fig 5a). These measured approximately 30 nm across, slightly more than half that reported by Moitra. At a higher concentration of 2.5 mg/mL, larger, spheroidal agglomerate structures developed. More notably, longer, fibrous structures also began appearing in the AFM scan (Fig 5b). In contrast to the spheroidal structures, these fibers appeared lower than the background. This apparent indentation is associated with formation of an aqueous film on the hydrophilic mica surface[24]. While this film conformed and added to hydrophilic surfaces, an opposite effect occurred with hydrophobic surfaces. The hydrophobicity of this peptide may be described as a function of structure. While the randomly packed spherical structures would produce a hydrophilic surface, akin to a micelle, the β-sheet structure associated with the nanotubes would produce relatively hydrophobic structure due to the phenyl side-chains. While these new fibrous structures were unique to the higher concentration scan, they were narrower than those previously reported. Nanotubes with diameters of 200 nm were expected; nanotubes with diameters of approximately 90 nm were measured. This smaller diameter was likely the result of a reduced peptide concentration, approximately half that demonstrated by Moitra. Lower peptide concentrations are associated with smaller aggregrates, as seen with the spherical structures in Fig 5.

**Figure 5.**
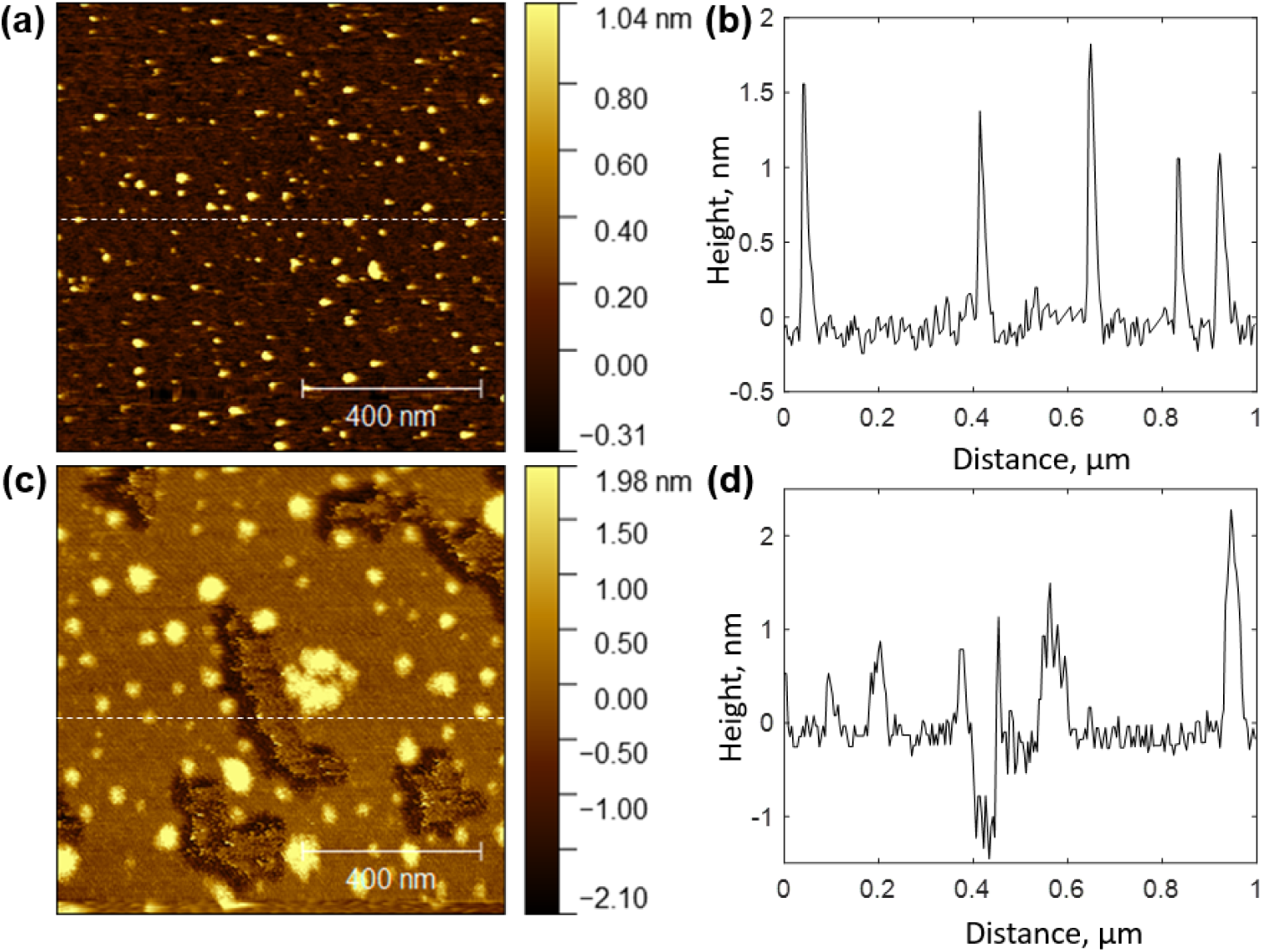
AFM scans of KFG tripeptide. **(a)** Scan of peptide deposited on mica at concentration of 0.5 mg/mL. **(b)** Topographical trace across dashed line shown in scan. **(c)** Scan of peptide deposited on mica at concentration of 2.5 mg/mL. **(d)** Topographical trace across dashed line shown in scan.

### Characterization of Fluorescent Peptide

A fluorescent hexapeptide was also demonstrated based on a prior work which studied the formation of a blue-emitting fluorophores as a result of photooxidation of specific tryptophan-containing peptides[25]. The hexapeptide was synthesized and oxidized under light in 2% hydrogen peroxide to produce a blue solution (Fig6a-b). With increasing exposure to this light, the intensity of blue fluorescence linearly increased until approximately 15 minutes, where emissions began to decrease (Fig 6c). This eventual decrease to signal is associated with degradation of the fluorescent structure by nonspecific oxidation. An initially high fluorescence intensity, slightly over half the maximum intensity, is associated with photooxidation in ambient room light during peptide synthesis, cleavage from the chlorotrityl resin, and purification (approximately 6 h).

**Figure 6.**
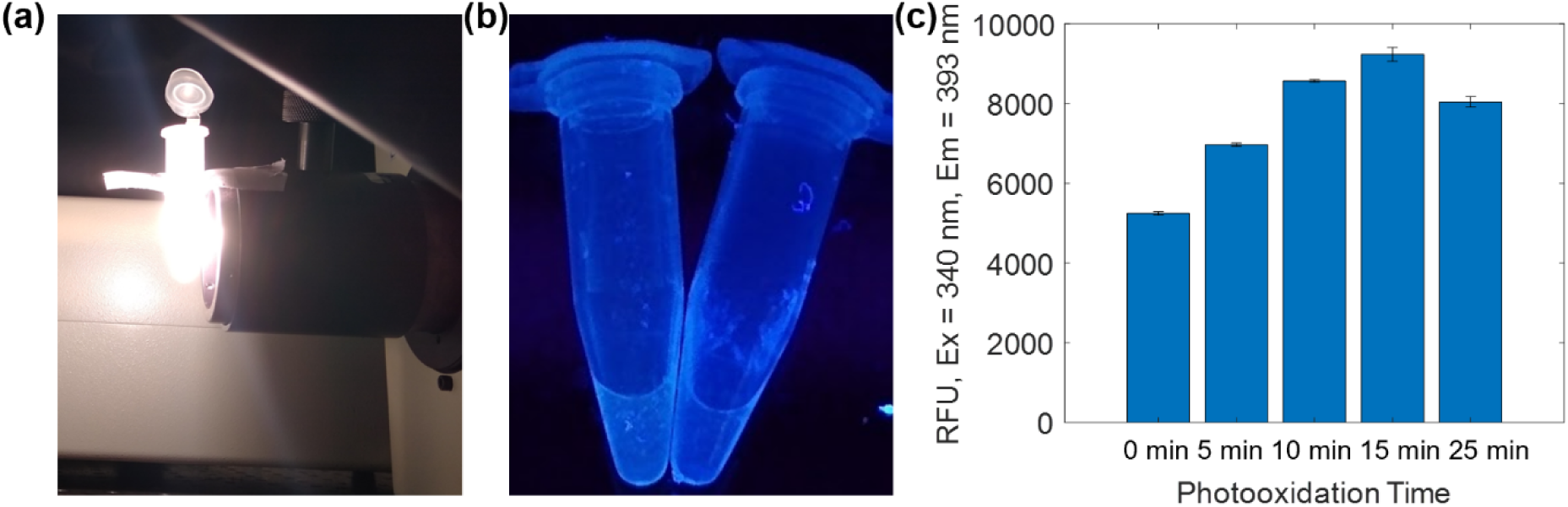
**(a)** Preparation of fluorescent peptide by photooxidation over high-power halogen lamp. **(b)** Fluorescent peptides over UV trans-illuminator. **(c)** Fluorescence intensities of peptides after exposure to lamp.

## Conclusion

In summary, we have demonstrated a prototype peptide synthesizer that utilizes inexpensive fluid-handling components to achieve a price point below US$300. While this prototype is only capable of handling 6 different amino acids in its current format, this design can be easily expanded to include 26 different amino acids while only raising the price to approximately US$600 (adding more pumps). We have used this prototype to synthesize several different peptides with different binding, structural, and chemical properties. With this peptide synthesizer, yields on the micromole scale can be achieved at costs of approximately $1/residue.

Although the price point of this synthesizer does make it available to a wider userbase, its use should be restricted to individuals with training in laboratory safety and access to fume hoods and proper chemical waste disposal (not a ‘do-it-yourself’ project for a garage). As with all self-built, electronic projects, precautions should be taken to minimize risk of fire in case of equipment malfunction, such as fire extinguisher training and minimization of combustibles. This work currently qualifies as a laboratory ‘frugal’ build, with some components having a price point still out of reach for some global researchers. However, further development of alternative, higher-volume parts for this design could reduce the price barrier and enable high-impact work in other limited-resource settings, thus democratizing access to peptides for new research.

This frugal synthesizer will accelerate the discovery process of novel peptide-based drugs and functionalization of nanomaterials for sensing and drug delivery purposes, particularly in laboratories lacking specialization in organic synthesis. Often there is a bottleneck in such groups (in terms of both funding and time) to obtain new peptide candidates from catalogues or from service vendors. While operating temperatures of this synthesizer can be increased to improve throughput, production of large (100+) peptide libraries may be better accomplished with more expensive, higher efficiency reactors capable of parallel synthesis. This frugal synthesizer can also function as an inexpensive educational tool for teaching labs (that have the sufficient safety measures in place, listed above), where low capital costs must be maintained. Moreover, it is a very useful pedagogical tool to have students build simple measurement systems.

## Acknowledgements

Work was supported in part by startup funds from Iowa State University, the Black & Veatch Building a World of Difference Faculty Fellowship in Engineering to NFR, the USDA Agricultural and Food Research Initiative Workforce and Education Development Program (Award # 2019-67011-29517) to NEK, and the Griswold undergraduate research internship to NER. The funders had no role in study design, data collection and analysis, decision to publish, or preparation of the manuscript.

## Notes

### Competing Interest Statement

The authors have declared no competing interest.

