## Supplemental Information for "Design and Validation of a Frugal, Automated, Solid-Phase Peptide Synthesizer"

### Supplement 1. Wiring Diagrams

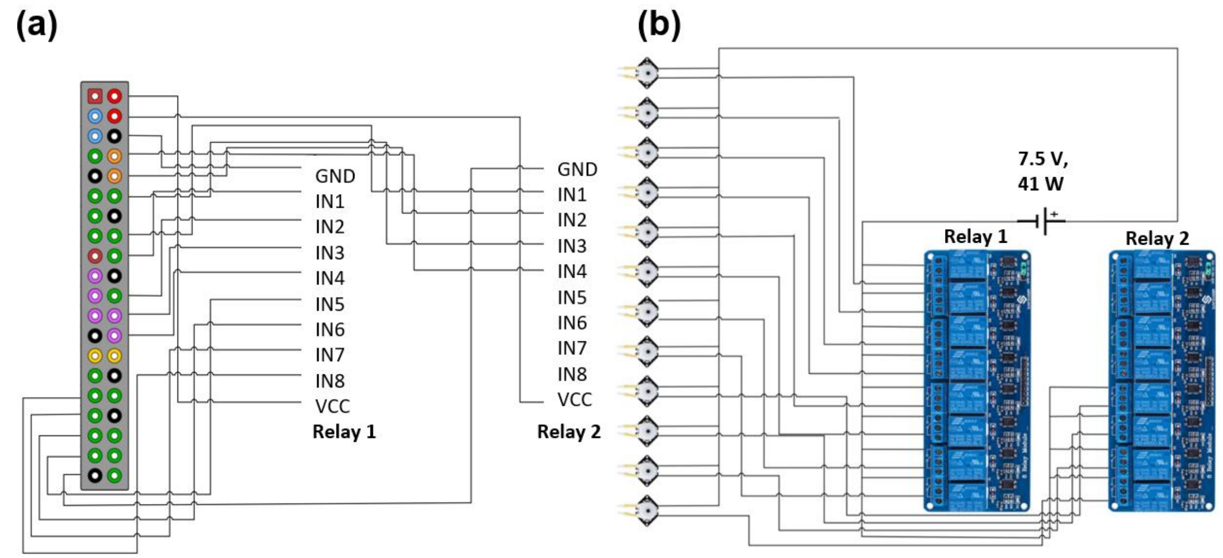

**Fig S1**. Wiring diagrams of device. (**a**) Connection of Raspberry Pi 3B GPIO pins to relays. (**b**) Connection of DC power supply to relays and pump motors.

### Supplement 2. Sources and Prices of Reagents

**Table S1**. Sources and prices of common SPPS reagents

| Reagent | Supplier | Quantity | Price (USD)<br>*Includes Hazmat packaging charge |
|---|---|---|---|
| NMM | Millipore Sigma | 250 mL | 22.70 |
| DMF | Chem-Impex | 1 L | 43.50* |
| HATU | Chem-Impex | 5 g | 36.00* |
| Piperidine | Chem-Impex | 1 L | 108.50* |
| Fmoc-glycine-2-chlorotrityl resin | Chem-Impex | 1 g | 20.00 |
| $N^{\varepsilon}$-Boc-L-lysine-2-chlorotrityl resin | Chem-Impex | 1 g | 20.00 |
| Aminomethyl polystyrene resin | Chem-Impex | 5 g | 20.00 |

**Table S2.** Quantities and prices of protected amino acids from ChemImpex

| Reagent | Quantity | Price (USD) |
|---|---|---|
| Na-Fmoc-Nw-(2,2,4,6,7-pentamethyldihydrobenzofuran-5-sulfonyl)-L-arginine | 5 g | 25.00 |

| | | |
|---|---|---|
| Fmoc-L-aspartic acid b-tert-butyl ester | 5 g | 12.50 |
| Fmoc-L-valine | 5 g | 10.00 |
| Fmoc-L-alanine | 5 g | 10.00 |
| Na-Fmoc-Nin-Boc-L-tryptophan | 5 g | 15.00 |
| Fmoc-O-tert-butyl-L-serine | 5 g | 15.00 |
| Na-Fmoc-Nim-trityl-L-histidine | 5 g | 15.00 |
| Fmoc-L-proline | 5 g | 10.00 |
| Na-Fmoc- Nd-trityl-L-glutamine | 5 g | 15.00 |
| Fmoc-L-phenylalanine | 5 g | 10.00 |
| Na-Fmoc-Ne-Boc-L-lysine | 5 g | 10.00 |
| Fmoc-L-glutamic acid g-tert butyl ester hydrate | 5 g | 15.00 |
| Fmoc-L-leucine | 5 g | 10.00 |
| Fmoc-O-tert-butyl-L-tyrosine | 5 g | 20.70 |
| Na-Fmoc-Nd-trityl-L-glutamine | 5 g | 15.00 |
| Fmoc-glycine | 25 g | 12.50 |
| Na-Fmoc-Ng-trityl-L-asparagine | 5 g | 12.50 |

### Supplement 3. Python3 Script for Synthesizer Operation

```
#!/usr/bin/env python
'''
Code for automated peptide synthesis
Step 1: Count the number of different amino acids used in your peptide.
Step 2: Connect GPIO pins from the Pi to the relay (use number of
different
        peptides plus 5 GPIO pins). Be sure to adjust the number of relay
        channels in the code as well.
Step 3: Pump 0 should pump pure DMF;
        Pump 1 should pump 20% piperidine in DMF;
        Pump 2 should pump the NMM and coupling agent;
        Pump 3 should be the draining pump for the bottom reaction vessel
        Pump 4 should be the draining pump for the top reaction vessel
Step 4: Starting from the end closest to the resin, use pump 5 for the
first
        amino acid, pump 6 for the second, etc. If an amino acid occurs
more
        than once, do not use another pump for the same amino acid.
Step 5: Run this Python file (make sure Python 3 is being used).
Step 6: Once the the file begins to run, type in the letter abbreviation
of the
        first peptide you will be using.
Step 7: The rest of the process should be done manually, but observe the
```

```
        process at all times to make sure everything runs smoothly.
'''
import RPi.GPIO as GPIO
from time import sleep
import numpy as np

'''
Initializes the GPIO pins utilized for the peptide synthesis.
'''
Relay_channel = [8,10,12,16,40,22,24,26,31,33,35,37]

'''
Initializes outputs for GPIO pins.
'''
def setup():
        GPIO.setmode(GPIO.BOARD)
        GPIO.setup(Relay_channel, GPIO.OUT, initial=GPIO.HIGH)

def main():
        '''
        Drains DMF manually added to reaction vessel once 10-15 minutes
        have been allotted for swelling.
        '''
        print('DMF from resin swelling being drained.')
        GPIO.output(Relay_channel[3], GPIO.LOW)
        sleep(20)
        GPIO.output(Relay_channel[3], GPIO.HIGH)
        sleep(1)

        '''
        Validates whether peptide the user entered is valid. If it is not,
        the user is prompted to enter the name of the peptide again.
        '''
        p = 'No'
        while p != 'Yes':
                AllAmiAci =
                ['A','R','N','D','B','C','Q','E','Z','G','H','I','L','K','
                M','F','P','S','T','W','Y','V']
                print('Please write out your peptide using single letter
abbreviations.')
                print('Be sure to use all capital letters.')
                Pep = input()
                t = 'F'
                while t == 'F':
                        if set(Pep) < set(AllAmiAci):
                                t = 'T'
                                PepLen = len(Pep)
                                NumAmiAci = len(set(Pep))
                        else:
                                t = 'F'
                                print('Invalid input. Please try again.')
                                print('Please write out your peptide using
                                      single letter abbreviations.')
                                print('Be sure to use all capital
```

```
letters.')
                                Pep = input()
                print('Is ',Pep, ' the amino acid chain you would like to
                      build (type \"Yes\" or \"No\")?')
                p = input()
                if p != 'Yes':
                        print('Since this is not the peptide you wanted or
                               you do not know how to type')
                        print('\"Yes\" properly, please start the process
over.')

        '''
        Ensures user is ready to begin the peptide synthesis. If they are
        not, prompts user to begin when they are ready.
        '''
        print('Are you ready to begin your peptide synthesis (type \"Yes\"
or \"No\")?')
        r = 'F'
        while r == 'F':
                y = input()
                if y != 'Yes':
                        print('Since you are not ready or you do not know
how to type')
                        print('\"Yes\" properly, type \"Yes\" when you are
ready.')
                        r = 'F'
                else:
                        r = 'T'

        '''
        Creates an array for each amino acid in the peptide corresponding
        to the position where it first appears in the sequence. For
        example, GSGS would correspond to 1212.
        '''
        for k in range(0,PepLen-1):
                PepNum = np.linspace(1,4,PepLen)
        for i in range(0,PepLen):
                j = 0
                q = 0
                while Pep[j] != Pep[i]:
                        j = j+1
                for c in range(1,j+1):
                        for z in range(0,c):
                                if Pep[c] == Pep[z]:
                                        q = q+1
                                        break
                PepNum[i] = int(round(j-q))

        '''Adds the current amino acid to the peptide chain. '''

        for l in range(0,PepLen):
                print('Amino acid',l+1,'which is',Pep[l],'and is in in
pump',str(PepNum[l]+5)[0])
                print('will now be added. Please standby.')
```

```
'''
Adds 20% piperidine in DMF to the solution two times (one
for two minutes and the second time for 6 minutes) to
deprotect the peptide that is being added.
'''
print('20% piperidine in DMF being added (1st time).')
GPIO.output(Relay_channel[1], GPIO.LOW)
sleep(0.6)
GPIO.output(Relay_channel[1], GPIO.HIGH)
sleep(120)
GPIO.output(Relay_channel[3], GPIO.LOW)
sleep(20)
GPIO.output(Relay_channel[3], GPIO.HIGH)
sleep(1)
print('20% piperidine in DMF being added (2nd time).')
GPIO.output(Relay_channel[1], GPIO.LOW)
sleep(0.6)
GPIO.output(Relay_channel[1], GPIO.HIGH)
sleep(480)
GPIO.output(Relay_channel[3], GPIO.LOW)
sleep(20)
GPIO.output(Relay_channel[3], GPIO.HIGH)
sleep(1)

'''
Washes the solution 4 times with DMF to rinse impurities
out of the reaction vessel.
'''
print('DMF washing steps now taking place.')
for k in range(0,4):
        GPIO.output(Relay_channel[0], GPIO.LOW)
        sleep(2)
        GPIO.output(Relay_channel[0], GPIO.HIGH)
        sleep(1)
        GPIO.output(Relay_channel[4], GPIO.LOW)
        sleep(20)
        GPIO.output(Relay_channel[4], GPIO.HIGH)
        sleep(1)
        GPIO.output(Relay_channel[3], GPIO.LOW)
        sleep(20)
        GPIO.output(Relay_channel[3], GPIO.HIGH)
        sleep(1)
'''
Adds a HATU solution dissolved in DMF and the amino acid
being added (also dissolved in DMF) to the reaction vessel
to add the current amino acid to the chain.
'''
print('Amino acid now being added.')
GPIO.output(Relay_channel[int(PepNum[l]+5)], GPIO.LOW)
sleep(1.25)
GPIO.output(Relay_channel[int(PepNum[l]+5)], GPIO.HIGH)
sleep(1)
GPIO.output(Relay_channel[2], GPIO.LOW)
```

```
                sleep(0.625)
                GPIO.output(Relay_channel[2], GPIO.HIGH)
                sleep(10)
                GPIO.output(Relay_channel[4], GPIO.LOW)
                sleep(20)
                GPIO.output(Relay_channel[4], GPIO.HIGH)
                sleep(900)
                GPIO.output(Relay_channel[3], GPIO.LOW)
                sleep(20)
                GPIO.output(Relay_channel[3], GPIO.HIGH)
                sleep(1)

                '''
                Washes the solution 4 times with DMF to rinse impurities
                out of the reaction vessel.
                '''
                print('DMF washing steps now taking place.')
                for k in range(0,3):
                        GPIO.output(Relay_channel[0], GPIO.LOW)
                        sleep(2)
                        GPIO.output(Relay_channel[0], GPIO.HIGH)
                        sleep(1)
                        GPIO.output(Relay_channel[4], GPIO.LOW)
                        sleep(20)
                        GPIO.output(Relay_channel[4], GPIO.HIGH)
                        sleep(1)
                        GPIO.output(Relay_channel[3], GPIO.LOW)
                        sleep(20)
                        GPIO.output(Relay_channel[3], GPIO.HIGH)
                        sleep(1)

                '''
                Notifies the user the current amino acid is done being
                added and asks if the user is ready to add the next amino
                acid in the sequence (if applicable). If the sequence
                completed, the code terminates.
                '''
                print('This amino acid step is complete.')
                if l != PepLen-1:
                        print('Are you ready to add your next amino acid
(type \"Yes\" or \"No\")?')
                        u = 'F'
                        while u == 'F':
                                p = input()
                                if p != 'Yes':
                                        print('Since you are not ready or
                                              you do not know how to
                                              type')
                                        print('\"Yes\" properly, type
\"Yes\" when you are ready.')
                                        u = 'F'
                                else:
                                        u = 'T'
                                        print('Moving on to the next amino
```

```
acid.')
                else:
                        print('Your peptide is done being made.')
                        print('Be sure to dispose of everything properly
and to clean all equipment.')

'''
Default settings if the code is terminated anytime during its execution.
'''
def destroy():
        GPIO.output(Relay_channel, GPIO.LOW)
        GPIO.cleanup()

if __name__ == '__main__':
        setup()
        try:
                main()
        except KeyboardInterrupt:
                destroy()
```

### Supplement 4. Reagent Pricing

**Table S-3**. SPPS Reagent Pricing

| **Reagent** | **Initially used** | **Amt per step** | **Cost/mL or g** | **Cost/Step, $** | **Initial Cost** |
|---|---|---|---|---|---|
| DMF | 100mL | 12.45mL | 0.0435 | 0.541575 | 0.435 |
| Piperidine | 0mL | 0.3mL | 0.1085 | 0.03255 | 0 |
| NMM | 0mL | 0.026875mL | 0.0908 | 0.00244025 | 0 |
| HATU | 0g | 0.046875g | 7.2 | 0.3375 | 0 |
| Amino acid | 0g | 0.0405g | 2.5 | 0.10125 | 0 |
| Resin | 0.05g | 0g | 4 | 0 | 0.2 |
| | | | **Startup Cost, $** | 0.635 | |
| | | | **Cost/Step, $** | 1.01531525 | |

### Supplement 5. Testing HA-tag peptide

Tests performed with HA-tags and anti-HA-tag monoclonal antibodies were less conclusive (Fig 3). While HA-tagged beads, on average, were slightly brighter than control beads, fragmented resin beads exhibited significantly brighter fluorescence. This may result from weaker HA-tag antibody affinity relative to streptavidin affinity and polystyrene-protein affinity.

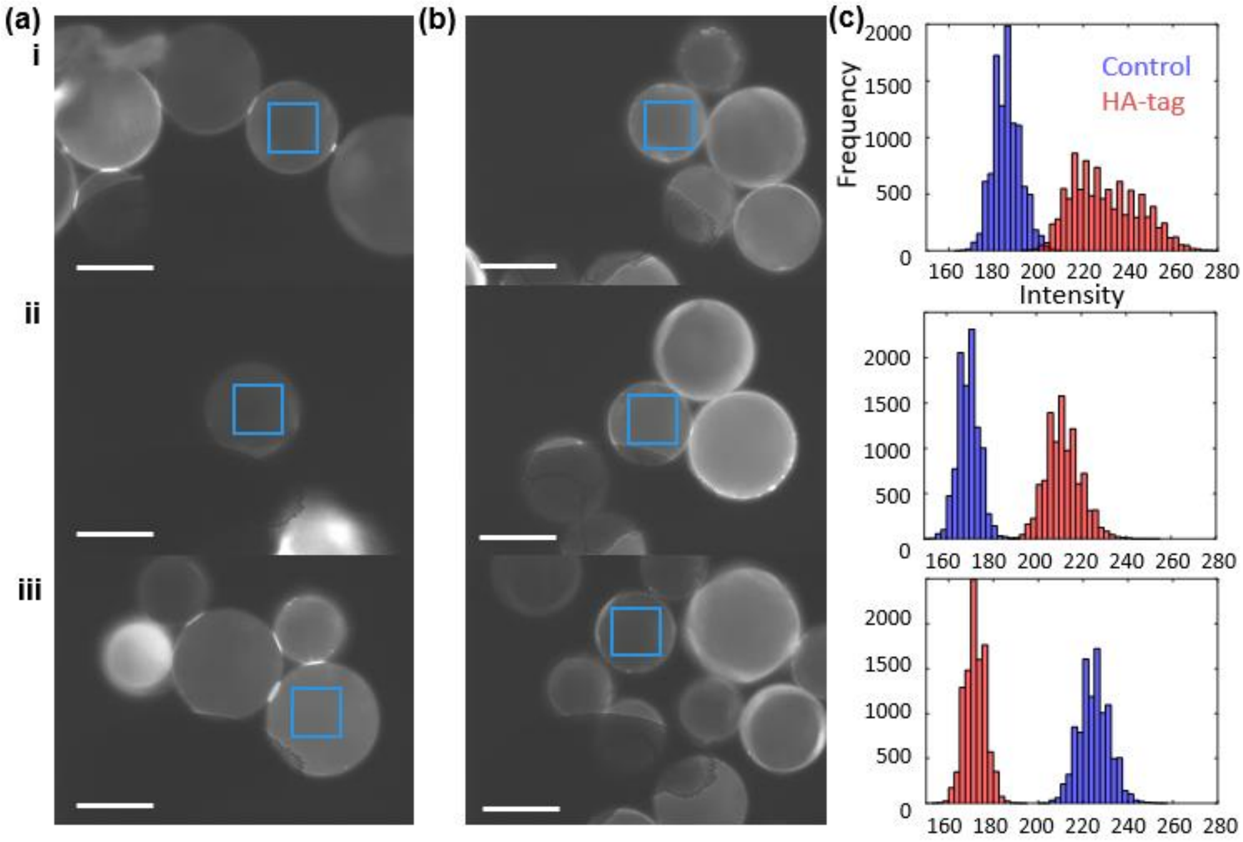

**Fig S-2.** Fluorescent microscope image of aminomethyl styrene resin functionalized with **(a)** control $(GS)_4$ sequence and **(b)** strep-tag after incubated with DyLight 488-anti-HA tag. **(c)** Histogram comparison of pixel intensities collected from regions of interest (solid blue boxes in a-b). **(i-iii)** Replicates. Scale bars are 50 µm.
